# *De novo* Genome Assembly of *Geosmithia morbida*, the Causal Agent of Thousand Cankers Disease

**DOI:** 10.1101/036285

**Authors:** Taruna Aggarwal, Anthony Westbrook, Kirk Broders, Keith Woeste, Matthew D MacManes

## Abstract

**Background**: Geosmithia morbida is a filamentous ascomycete that causes Thousand Cankers Disease in the eastern black walnut tree. This pathogen is commonly found in the western U.S.; however, recently the disease was also detected in several eastern states where the black walnut lumber industry is concentrated. G. morbida is one of two known phytopathogens within the genus Geosmithia, and it is vectored into the host tree via the walnut twig beetle.

**Results**: We present the first de novo draft genome of G. morbida. It is 26.5 Mbp in length and contains less than 1% repetitive elements. The genome possesses an estimated 6,273 genes, 277 of which are predicted to encode proteins with unknown functions. Approximately 31.5% of the proteins in G. morbida are homologous to proteins involved in pathogenicity, and 5.6% of the proteins contain signal peptides that indicate these proteins are secreted.

**Conclusions**: Several studies have investigated the evolution of pathogenicity in pathogens of agricultural crops; forest fungal pathogens are often neglected because research efforts are focused on food crops. G. morbida is one of the few tree phytopathogens to be sequenced, assembled and annotated. The first draft genome of G. morbida serves as a valuable tool for comprehending the underlying molecular and evolutionary mechanisms behind pathogenesis within the Geosmithia genus.

## Introduction

Studying molecular evolution of any phenotype is now made possible by the analysis of large amounts of sequence data generated by next-generation sequencing platforms. This is particularly beneficial in the case of emerging fungal pathogens, which are progressively recognized as a threat to global biodiversity and food security. Furthermore, in many cases their expansion is a result of anthropogenic activities and an increase in trade of fungal-infected goods [1]. Fungal pathogens evolve in order to overcome host resistance, fungicides, and to adapt to new hosts and environments. Whole genome sequence data have been used to identify the mechanisms of adaptive evolution within fungi [2–4]. For instance, Stukenbrock et al. (2011) investigated the patterns of evolution in fungal pathogens during the process of domestication in wheat using all aligned genes within the genomes of wheat pathogens. They found that *Mycosphaerella graminicola,* a domesticated wheat pathogen (now known as *Zymoseptoria tritici),* underwent adaptive evolution at a higher rate than its wild relatives, *Mycosphaerella* S1 and *Mycosphaerella* S2. The study also revealed that many of the pathogen's 802 secreted proteins were under positive selection. A study by Gardiner et al. (2012), identified genes encoding aminotransferases, hydrolases, and kinases that were shared between *Fusarium pseudograminearum* and other cereal pathogens. Using genomic and phylogenetic analyses, the researchers demonstrated that these genes had bacterial origins. These studies highlight the various evolutionary means that fungal species employ in order to adapt to specific hosts, as well as the important role genomics and bioinformatics play in elucidating evolutionary mechanisms within the fungal kingdom.

Many tree fungal pathogens associate with bark beetles, which belong to the Scolytinae family [5]. As climate patterns change, both the beetles and their fungal symbionts are able to invade new territory and become major invasive forest pests on a global scale [6, 7]. A well-known example of an invasive pest is the mountain pine beetle and its symbiont, *Grosmannia clavigera* that has affected approximately 3.4 million of acres of lodgepole, ponderosa, and five-needle pine trees in Colorado alone since the outbreak began in 1996 [8, 9]. Another beetle pest in the western U.S., *Pityophthorous judlandis* (walnut twig beetle), associates with several fungal species, including the emergent fungal pathogen *Geosmithia morbida* [10].

Reports of tree mortality triggered by *G. morbida* infections first surfaced in 2009 [12], and the fungus was described as a new species in 2011 [10]. This fungus is vectored into the host via *P. juglandis* and is the causal agent of thousand cankers disease (TCD) in *Julgans nigra* (eastern black walnut) [12]. This walnut species is valued for its wood, which is used for furniture, cabinetry, and veneer. Although *J. nigra* trees are planted throughout western U.S. as a decorative species, they are indigenous to eastern North America where the walnut industry is worth hundreds of millions of dollars [13]. In addition to being a major threat to the eastern populations of *J. nigra,* TCD is of great concern because certain western walnut species including *J. regia* (the Persian walnut), *J. californica,* and *J. hindsii* are also susceptible to the fungus according to greenhouse inoculation studies [14].

The etiology of TCD is complex because it is a consequence of a fungal-beetle symbiosis. The walnut twig beetle, which is only known to attack members of genus *Juglans* and *Pterocarya,* is the most common vector of *G. morbida* [10]. Nevertheless, other beetles are able to disperse the fungus from infested trees [15, 16]. As vast numbers of beetles concentrate in the bark of infested trees, fungal cankers form and coalesce around beetle galleries and entrance holes. As the infection progresses, the phloem and cambium discolor and the leaves wilt and yellow. These symptoms are followed by branch dieback and eventual tree death, which can occur within three years of the initial infection [10]. Currently, 15 states in the U.S. have reported one or more incidences of TCD, reflecting the expansion of WTB's geographic range from its presumed native range in a few southwestern states [17].

To date, *G. morbida* is one of only two known pathogens within the genus *Geosmithia,* which consists of mostly saprotrophic beetle-associated species (the other pathogen is *G. pallida)* [18]. The ecological complexity this vector-host-pathogen complex exhibits makes it an intriguing lens for studying the evolution of pathogenicity within the fungal kingdom. A well-assembled reference genome will enable us to identify genes unique to *G. morbida* that may be utilized to develop sequence-based tools for detecting and monitoring epidemics of TCD and for studying the evolution of pathogenesis within the *Geosmithia* genus. Here, we present a *de novo* genome assembly of *Geosmithia morbida.* The objectives of this study are to: 1) assemble the first, high-quality draft genome of this pathogen; 2) annotate the genome in order to better comprehend the evolution of pathogenicity in the *Geosmithia* genus; and 3) briefly compare the genome of *G. morbida* to two other fungal pathogens for which genomic data is available: *Fusarium solani,* a root pathogen that infects soybean, and *Grosmannia calvigera,* a pathogenic ascomycete that associates with the mountain pine beetle and kills lodgepole pines in North America.

## Methods

### DNA extraction and Library Preparation

DNA was extracted using the CTAB method as outlined by the Joint Genome Institute to extract DNA for Genome Sequencing from lyophilized mycelium of *G. morbida* (isolate 1262, host: *Juglans californica)* from southwestern California [19]. The total DNA concentration was measured using Nanodrop, and samples for sequencing were sent to Purdue University Genomics Core Facility in West Lafayette, Indiana. DNA libraries were prepared using the paired-end Illumina Truseq protocol and mate-pair Nextera DNA Sample Preparation kits with average insert sizes of 487bp and 1921bp respectively. These libraries were sequenced on the Illumina HiSeq 2500.

### Preprocessing Sequence Data

We began by performing quality control checks on our raw sequence data generated by the Illumina platform. To assess the quality of our data, we ran FastQC (v0.11.2) (https://goo.gl/xHM1zf) [20] and SGA Preqc (v0.10.13) (https://goo.gl/9y5bNy) on our raw sequence reads [21]. Both tools aim to supply the user with information such as per base sequence quality score distribution (FastQC) and frequency of variant branches in *de Bruijn* graphs (Preqc) that aid in selecting appropriate assembly tools and parameters. The paired-end raw reads were corrected using a Bloom filter-based error correction tool called BLESS (v0.16) (https://goo.gl/Kno6Xo) [22]. Next, the error corrected reads were trimmed with Trimmomatic, version 0.32, using a Phred threshold of 2, following recommendations from MacManes (2014) (https://goo.gl/FFoFjL) [23]. NextClip, version 1.3.1, was leveraged to trim adapters in the mate-pair read set (https://goo.gl/aZ9ucT) [24]. The raw reads are available at https://goo.gl/IMsMe5.

### *De novo* genome assembly and evaluation

The *de novo* genome assembly was constructed with ALLPaths-LG (v49414) (https://goo.gl/03gU9Z) [25]. The assembly was evaluated with BUSCO (v1.1b1) (https://goo.gl/bMrXIM), a tool that assesses genome completeness based on the presence of single-copy orthologs [26]. We also generated length-based statistics for our *de novo* genome with QUAST (v2.3) (https://goo.gl/5KSa4M) [27]. The raw reads were mapped back to the genome using BWA version 0.7.9a-r786 to further assess the quality of the assembly (https://goo.gl/Scxgn4) [28]

### Structural and Functional Annotation of *G. morbida* genome

We used the automated genome annotation software Maker version 2.31.8. Maker identifies repetitive elements, aligns ESTs, and uses protein homology evidence to generate *ab initio* gene predictions (https://goo.gl/JiLA3H) [29]. We used two of the three gene prediction tools available within the pipeline, SNAP and Augustus. SNAP was trained using gff files generated by CEGMA v2.5 (a program similar to BUSCO). Augustus was trained with *Fusarium solani* protein models (v2.0.26) downloaded from Ensembl Fungi [30, 34]. In order to functionally annotate the genome, the protein sequences produced by the structural annotation were blasted against the Swiss-Prot database, and target sequences were filtered for the best hits [31]. A small subset of the resulting annotations was visualized and manually curated in WebApollo v2.0.1 [32]. The final annotations were also evaluated with BUSCO (v1.1b1) (https://goo.gl/thTGzH).

### Assessing Repetitive Elements Profile

To assess the repetitive elements profile of *G. morbida,* we masked only the interspersed repeats within the assembled scaffolds with RepeatMasker (v4.0.5) (https://goo.gl/TXrbr3) [33] using the sensitive mode and default values as arguments. In order to compare the repetitive element profile of *G. morbida* with *F. solani* (v2.0.29) and *G. clavigera* (kw1407.GCA_000143105.2.30), the interspersed repeats of these two fungal pathogens were also masked with RepeatMasker. The genome and protein data of these fungi were downloaded from Ensembl Fungi [34].

### Identifying putative proteins contributing to pathogenicity

To identify putative genes contributing to pathogenicity in *G. morbida,* a BLASTp search was conducted for single best hits at an e-value threshold of 1e-6 or less against the PHI-base database (v3.8) (https://goo.gl/CEEVY0) that contains experimentally confirmed genes from fungal, oomycete and bacterial pathogens [35]. The search was performed using the same parameters for *F. solani* and *G. clavigera.* To identify the proteins that contain signal peptides, we used SignalP (v4.1) (https://goo.gl/JOe5Dh), and compared results from *G. morbida* with those from *F. solani* and G. *claviger* a [36]. Lastly, to find putative protein domains involved in pathogenicity in *G. morbida,* we performed a HMMER (version 3.1b2) [37] search against the Pfam database (v28.0) [38] using the protein sequences as query. We conducted the same search for sequences of 17 known effector proteins, then extracted and analyzed domains common between the effector sequences and *G. morbida* (https://goo.gl/Y9IPZs).

## Results and Discussion

### Data Processing

A total of 28,027,726 PE and 41,348,578 MP forward and reverse reads were generated with approximately 56x and 83x coverage respectively (Table 1). Of the MP reads, 67.7% contained adapters that were trimmed using NextClip (v1.3.1). We corrected errors within the PE reads using BLESS (v0.16) at a kmer length of 21. After correction, low-quality reads (phred score < 2) were trimmed with Trimmomatic (v0.32) resulting in 99.75% reads passing. In total, 16,336,158 MP and 27,957,268 PE reads were used to construct the *de novo* genome assembly.

**Table 1.** Statistics for *Geosmithia morbida* sequence data.

|  | Paired-end |  | Mate-pair |  |
| --- | --- | --- | --- | --- |
| Number of reads | 28,027,726 | **27,957,268** | 41,348,578 | **16,336,158** |
| Average insert size (bp) | 487 |  | 1921 |  |
| Average coverage | 56x |  | 83x |  |

The values in bold are number of trimmed, error corrected and filtered reads that were used for the assembly.

## Assembly Features

The *G. morbida de novo* assembly (available at https://goo.gl/6P8zmY) was constructed with AllPaths-LG (v49414). The assembled genome consisted of 73 contigs totaling 26,549,069 bp. The largest contig length was 2,597,956 bp, and the NG50 was 1,305,468 bp. The completeness of the genome assembly was assessed using BUSCO, a tool that scans the genome for the presence of single-copy orthologous groups present in more than 90% of fungal species. Of 1,438 single-copy orthologs specific to fungi, 95% were complete in our assembly, and 3.6% were fragmented BUSCOs. Only 0.8% of the orthologs were missing from the genome (Table 2). We used BWA to map the unprocessed, raw MP and PE reads back to the genome to further evaluate the assembly, and 87% of the MP and 90% of the PE reads mapped to our reference genome.

**Table 2.** *Geosmithia morbida* reference genome assembly statistics generated using QUAST (v2.3)

|  | Scaffolds |
| --- | --- |
| Number of sequences | 73 |
| Largest scaffold length | 2,597,956 |
| N50 | 1,305,468 |
| L50 | 7 |
| Total assembly length | 26,549,069 |
| GC% | 54.31 |
| BUSCOs completeness | 95% |

### Gene annotation

The automated genome annotation software Maker v2.31.8 was used to identify structural elements in the *G. morbida* assembly generated by AllPaths-LG. Of the total 6,273 proteins that were predicted, 5,996 returned with hits against the Swiss-Prot database–only 277 (4.41%) of the total genes encoded for proteins of unknown function. The completeness of the functional annotations was evaluated using BUSCO, and 94% of the single copy orthologs were present in this protein set. The transcript and protein files are available at https://goo.gl/svTmKp and https://goo.gl/pB9y5l.

### Repetitive Elements

Repetitive elements represented 0.81% of the total bases in the *G. morbida* genome (available at https://goo.gl/wDq2xP). The genome contained 152 retroelements (class I) that were mostly composed of long terminal repeats (n=146) and 60 DNA transposons (class II). In comparison, the genomes of *G. clavigera* and *F. solani* contained 1.14% and 1.47% respectively (available at https://goo.gl/8zXAIH and https://goo.gl/YQAM2N). *G. clavigera* possesses 541 retroelements (0.79%) and 66 DNA transposons (0.04%), whereas the genome of *F. solani* is comprised of 499 (0.54%) and 515 (0.81%) retroelements and transposons respectively. The larger number of repeat elements in *F. solani* may explain its relatively large genome size −51.3 Mbp versus *G. clavigera's* 29.8 Mbp and *G. morbida's* 26.5 Mbp (Table 3).

**Table 3.**
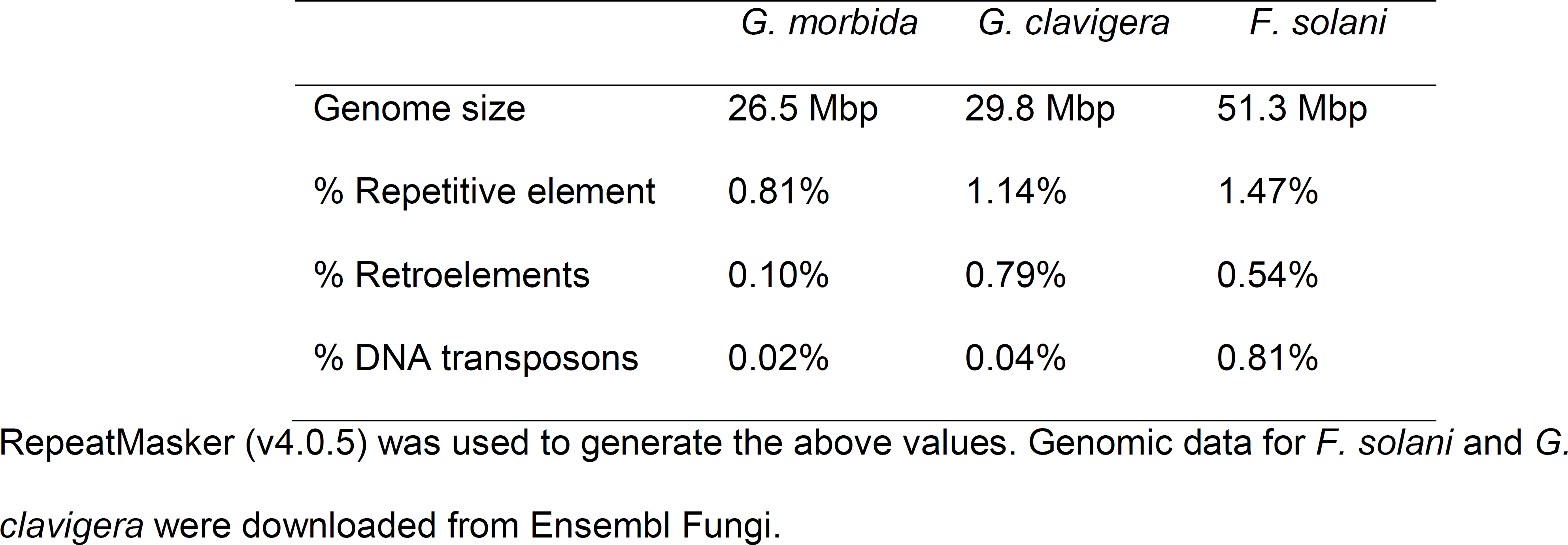
Repetitive elements profile for *Geosmithia morbida, Grosmannia clavigera* and *Fusarium solani.*

## Identifying and classifying putative pathogenicity genes

We blasted the entire predicted protein set against the PHI-base database (v3.8) to identify a list of putative genes that may contribute to pathogenicity within *G. morbida, F. solani,* and *G. clavigera.* We determined that 1,974 genes in *G. morbida* (31.47% of the total 6,273 genes) were homologous to protein sequences in the database (available at https://goo.gl/SZA4Kd). For *F. solani* and *G. clavigera,* there were 4,855 and 2,387 genes with homologous PHI-base proteins (available at https://goo.gl/Rm8Zx7 and https://goo.gl/fjrrvm).

### Identifying putative secreted proteins

A search for the presence of secreted peptides within the protein sequences of *G. morbida, F. solani* and *G. clavigera* showed that approximately 5.6% (349) of the *G. morbida* protein sequences contained putative signal peptides (available at https://goo.gl/Qz8gUr). Of the 349 sequences containing putative signal peptides, only 27 encoded proteins of unknown function. Roughly 8.8% and 6.9% of the proteins of *F. solani* and *G. clavigera* possess signal peptides (available at https://goo.gl/mTu7Ok and https://goo.gl/PZdSNc). Secreted proteins are essential for host-fungal interactions and are indicative of adaptation within fungal pathogens that require an array of mechanisms to overcome plant host defenses.

### Identifying protein domains

We conducted a HMMER search against the pfam database (v28.0) using amino acid sequences for *G. morbida* and 17 effector proteins from various fungal species. For *G. morbida,* there were 6,023 unique protein domains out of a total of 43,823 Pfam hits. A total of 17 domains, which comprised 1,000 hits, were shared between *G. morbida* and known effector proteins. The three most common protein domains in *G. morbida* with a putative effector function belonged to short-chain dehydrogenases (n=111), polyketide synthases (n=94) and NADH dehydrogenases (n=86). The HMMER *G. morbida* and effector proteins output files are located at https://goo.gl/r8B7uk and https://goo.gl/mkn5aB respectively.

## Conclusion

This work introduces the first genome assembly and analysis of *Geosmithia morbida,* a fungal pathogen of the black walnut tree that is vectored into the host via the walnut twig beetle. The *de novo* assembly is composed of 73 scaffolds totaling in 26.5 Mbp. There are 6,273 predicted proteins, and 4.41% of these are unknown. In comparison, 68.27% of *F. solani* and 26.70% of *G. clavigera* predicted proteins are unknown. We assessed the quality of our genome assembly and the predicted protein set using BUSCO, and found that 95% and 94% of the single copy orthologs specific to the fungal lineage were present in both respectively. These data are indicative of our assembly's high quality and completeness. Our BLASTp search against the PHI-base database revealed that *G. morbida* possesses 1,974 genes that are homologous to proteins involved in pathogenicity. Furthermore, *G. morbida* shares several domains with known effector proteins that are key for fungal pathogens during the infection process.

*Geosmithia morbida* is one of only two known fungal pathogens within the *Geosmithia* genus [18]. The genome assembly introduced in this study can be leveraged to explore the molecular mechanisms behind pathogenesis within this genus. The putative list of pathogenicity genes provided in this study can be used for future comparative genomic analyses, knock-out, and inoculation experiments. Moreover, genes unique to *G. morbida* may be utilized to develop DNA sequence-based tools for detecting and monitoring ongoing and future TCD epidemics.

## Authors' contributions

TA extracted DNA, prepared samples for sequencing, wrote the manuscript. TA and MDM assembled and evaluated the genome. AW annotated the genome. KB and KW helped conceive and fund the project and assisted in manuscript editing. MDM and KB contributed to the writing of the manuscript.

## Acknowledgements

Partial funding was provided by the New Hampshire Agricultural Experiment Station. This is Scientific Contribution Number 2652. Funding was also provided by the USDA Forest Service, Forest Health and Protection. Mention of a trademark, proprietary product, or vendor does not constitute a guarantee or warranty of the product by the U.S. Department of Agriculture and does not imply its approval to the exclusion of other products or vendors that also may be suitable.

